# Primary cortical neurons precipitate and extrude large mitochondria-associated calcium-phosphate sheets with a bone-precursor-like ultrastructure

**DOI:** 10.1101/2025.08.04.668590

**Authors:** Erik D. Anderson, Christopher A. Cronkite, Philip R. Baldwin, Carlota P. Abella, Joseph G. Duman, Ashleigh N. Simmonds, M. Neal Waxham, Kimberley F. Tolias, Steven J. Ludtke

**Affiliations:** Department of Biochemistry and Molecular Pharmacology, Baylor College of Medicine, Houston, TX, USA; Department of Neuroscience, Baylor College of Medicine, Houston, TX, USA; Department of Neurobiology and Anatomy, McGovern Medical School, University of Texas Health Science Center at Houston, Houston, TX, USA

**Keywords:** Calcium-phosphate, Neurons, Mitochondria, Migrasomes, Cryo-ET, LDSAED, Cryo-CLEM

## Abstract

Calcium-phosphate (CaP) is a ubiquitous inorganic compound that plays an important structural role in healthy bone and teeth formation, but its pathologic buildup can occur in dyshomeostatic calcium disorders like Alzheimer’s disease and Leigh syndrome. The nexus of pathologic extracellular CaP in the nervous system is not well understood, but prior evidence suggests mitochondria could be a source. We have observed mitochondria-sized sheet-like CaP aggregates within functional wild type cortical neuron cultures at 1 and 20 DIV. Neurons were extracted from embryonic day 18 (E18) rat embryos following standard protocols to study neuronal structure and function. We have used a combination of cryo-ET, cryo-CLEM, and LDSAED to demonstrate that these aggregates are octacalcium phosphate-like, are associated with mitochondria, and that at least a portion are extruded via migrasomes. Visually similar aggregates were previously observed in Huntington’s disease model neurons, but in that study they were not observed in WT controls. These findings show that this CaP aggregation process occurs routinely in WT neurons and may reveal an important link for how mitochondria may participate in calcification, highlighting them as potential therapeutic targets in neurological disorders characterized by pathological calcification, such as Alzheimer’s disease.

## Introduction

Calcium-phosphate (CaP) biomineralization is an important process that forms the inorganic component of bone and teeth. However, its deposition in the nervous system is also a prevalent symptom of brain pathology. It is seen in a wide variety of both rare and common diseases from inherited mitochondrial encephalopathies such as Leigh Syndrome to neurodegenerative disorders like Alzheimer’s disease (Tsolaki et al., 2022; Monfrini et al., 2023). CaP buildup in the brain can cause seizures [3], but where exactly this CaP originates is an open and important question.

Although physiologic biomineralization processes such as bone formation offer valuable insights, the underlying sequence of events involved remains incompletely understood and is subject to ongoing debate. Increasing evidence supports bone formation beginning with an amorphous calcium-phosphate (ACP, Ca_9_(PO_4_)_6_) precursor that lacks long range structure, which either progresses through an octacalcium phosphate-like (OCP, Ca_8_(HPO_4_)_2_(PO_4_)_4_·5H_2_O) intermediate or directly converts into hydroxyapatite (HA, Ca_10_(PO_4_)_6_(OH)_2_) [4–7]). This process has been shown to occur in extracellular matrix vesicles released from osteoblasts [8], but where the ACP originates is still debated. Molecules such as Annexin-related proteins [9] and poly(ADP-ribose) (Genge et al., 1990; Müller et al., 2019) that are enriched in matrix vesicles are thought to play an important role in calcium sequestration. However, the ACP granules that form in Ca_2_^2+^ and PO_4_^3-^ supersaturated mitochondria [12] have also been hypothesized to contribute CaP through mitophagy-dependent transfer of these granules to autolysosomes [13,14].

Anytime there are large, rapid calcium fluctuations in a cell, CaP precipitation is possible [15]. In neutral to slightly alkaline environments like the mitochondrial matrix, when calcium and phosphate ions reach supersaturation, ACP is frequently the first precipitate to form [16]. High levels of magnesium, citrate, and other ions in the matrix inhibit hydroxyapatite conversion (Duvvuri and Lood, 2021; Brdiczka and Barnard, 1979), however it has been reported in the mitochondria of mouse neurons following excitotoxic shock [18]. Hydroxyapatite has also been reported in the mitochondria of cardiomyocytes that have undergone ischemia-reperfusion injury, which occurs when blood flow is restored to previously oxygen-deprived tissue [19].

Ischemia causes mitochondrial calcium, ADP, and pyrophosphate concentrations to increase and pH to drop through decreased oxidative phosphorylation and increased glycolysis [20]. Following reperfusion, increased oxidative phosphorylation causes mitochondrial pH to rise. However, the concomitant high calcium concentration can induce the mitochondrial permeability transition pore (mPTP) to open, allowing calcium and other solutes <1.5 kDa to flow out of the matrix [20]. Classically, severe metabolic disturbances trigger programmed cell death via apoptosis or necrosis. In contrast, less severe ionic and oxygen-related stress leads to mitophagy, the partial or complete engulfment of damaged mitochondria into autophagosomes [21]. Interestingly an alternative mechanism for mitochondria quality control has recently been identified: some damaged mitochondria are actively extruded from cells in large (∼3 µm) vesicles called migrasomes, through a process termed mitocytosis [22]. Migrasomes are released on the lagging edge of migrating cells and have been identified in primary mouse hippocampal neurons (Ma et al., 2015, 2023). They are distinct from exosomes, and some were shown to rupture and release their contents into the extracellular space [24], providing a mechanism for how damaged mitochondria and their fragments can be found outside of cells.

Similar reperfusion-like injuries are difficult to avoid when isolating embryonic neurons for cell culture. A standard procedure that our lab follows to isolate rat cortical neurons uses a base media of ∼4° C Hanks’ balanced salt solution (HBSS) during benchtop dissection and ∼37° C Neurobasal media during trituration and subsequent culture [25,26]. Low temperature and no calcium during dissection helps prevent ischemic damage following removal from maternal circulation by slowing oxidative phosphorylation, limiting NMDA receptor activation and keeping the mPTP closed [27,28]. However, upon transfer into warm Neurobasal media with a higher calcium concentration of 1.8 mM, this ion will enter the cell, providing a setting where ACP could precipitate.

Recently, large unidentified sheet aggregates were observed in Huntington’s disease model neuronal cultures that were absent in WT control neuron cultures prepared under the same conditions [29]. These aggregates had a similar hyper-electron dense appearance to the ACP granules when visualized with cryo-electron tomography (cryo-ET), but with no apparent periodic structure. While the authors hypothesized that these aggregates might be within autophagosomes or mitophagy-related vesicles and were possibly derived from damaged mitochondria, this was not confirmed. The observed aggregates’ molecular constituents remained unknown, and they have not been reported in normal healthy neurons.

In this study we report mitochondria-sized sheet aggregates in WT rat embryonic neuron cultures at both 1 and 20 days *in vitro* (DIV) with an appearance similar to those reported in Huntington’s disease model neurons. In this study, we use a combination of cryo-ET, cryo-correlative light and electron microscopy (cryo-CLEM), and low-dose selected area electron diffraction (LDSAED) to characterize the aggregates. Our results suggest these aggregates are CaP precipitated in mitochondria which are then extruded at least in part by migrasomes. These results provide a mechanism for how mitochondria of cortical neurons may serve as a nexus for CaP precipitates in the central nervous system.

## Results

We isolated cortical neurons from E18 rat embryos and grew them on cryo-EM grids for 24 hours under conditions designed to optimize healthy growth. At this stage, viable neurons can be distinguished from non-viable neurons by the presence of extensive cytoplasm and the early formation of neurites (Fig. 1 A.1). In these cultures, we observe aggregates intracellularly (Fig. 1 A.2, A.4) in both cell bodies (Fig. 1 A.2) and neurites (Fig. 1 A.4) as well as extracellularly (Fig. 1 A.3). They appear to be in most neurons, though the number varies widely without any morphologic predictability (i.e., there are some neurons with abundant cytoplasm that have numerous aggregates, while others possess few or no aggregates). The same is true for neurons whose cytoplasm has retracted. Generally, there appear to be more aggregates in the soma than in neurites but calculating the abundance of aggregates/neuron or aggregates/cellular region is limited by the detection method. We chose cryo-transmission electron microscopic methods over room-temperature because cells are vitrified in a few milliseconds without necessary fixatives or stains that could affect the aggregate formation and structure. This may be why before Wu *et al.*’s 2023 study they were not reported. High resolution cryo-EM is currently the only method to identify them, but because it utilizes transmitted electrons, the thick center of a neuron’s soma makes it impossible to definitively know the total number in that area. While cryo-FIB/SEM can be used to thin these regions, the material above and below the section is lost in the process. Aggregates appear in a clear majority of neurons imaged and are more prevalent in the soma and extracellularly than in neurites.

**Figure 1.**
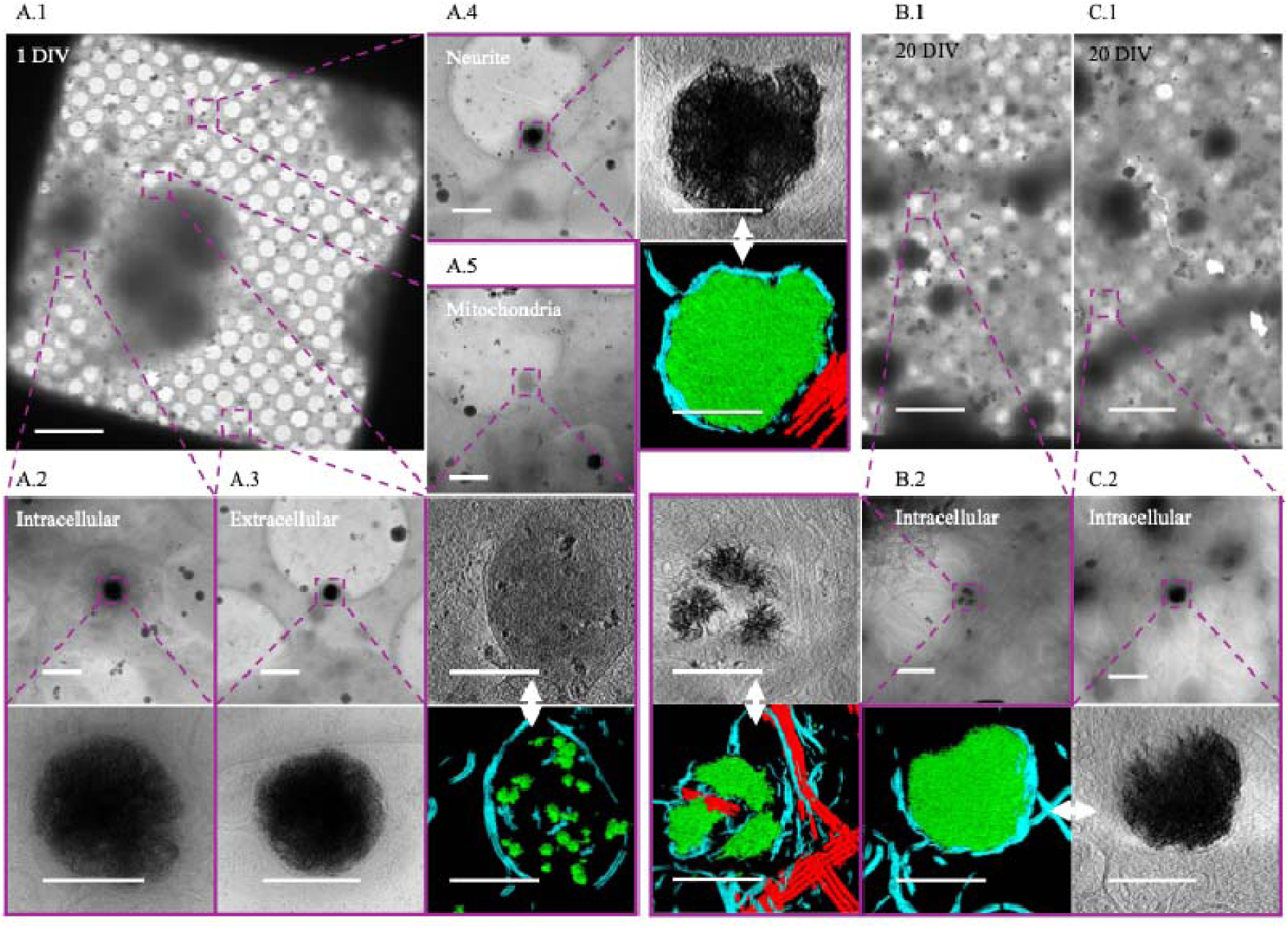
Sheet aggregates are present in both 1 DIV and 20 DIV rat cortical neurons. **(A.1)** 700× cryo-EM micrograph of a holey-carbon grid square depicting a typical vitrified cultured 1 DIV rat cortical neuron. **(A.2)** 5,300× (top) and high (bottom) mag cryo-EM micrograph images depicting aggregates within a soma. **(A.3)** 5,300× (top) and 45,000× (bottom) cryo-EM micrograph images showing an extracellular aggregate. **(A.4)** 5,300× cryo-EM micrograph (top left), 45,000× cryo-ET tomogram (top right), and 45,000× cryo-ET tomogram segmentation (bottom right) depicting aggregates within a neurite. **(A.5)** 5,300× cryo-EM micrograph (top), 45,000× cryo-ET tomogram (middle), and 45,000× tomogram segmentation (bottom) depicting mitochondria with calcium phosphate granules. **(B.1)** and **(C.1)** depict 700× cryo-EM micrographs of a typical vitrified cultured 20 DIV rat cortical neuron. **(B.2)** 5,300× cryo-EM micrograph (top right), 45,000× cryo-ET tomogram (top left), and 45,000× tomogram segmentation (bottom left) depicting aggregates. **(C.2)** 5,300× cryo-EM micrograph (top right), 45,000× cryo-ET tomogram (bottom right), and 45,000× tomogram segmentation (bottom left) depicting aggregates. Dashed boxes and lines depict areas of interest that are gradually increased in magnification. In segmentations, green = aggregates, cyan = membranes, and red = microtubules. Double-headed arrows depict corresponding tomogram and segmentation. Scale bars for 700× images = 10 µm; 5,300× = 1 µm; and 45,000× = 200 nm.

When viewed in 2D, as noted by Wu *et al*, the aggregates appear like fibers. However, with tomographic reconstruction and segmentation in 3D, it becomes clear that these apparent fibers are actually sheets (Fig. 2 A). The central region of more developed aggregates is generally amorphous in appearance and difficult to segment (Fig. 1 A.4). Aggregates such as these appear similar to the small calcium phosphate granules known to form in mitochondria (Fig. 1 A.3), though roughly 10x larger (sheet aggregates = ∼ 200 nm – > 1 µm; granules = ∼ 20 nm – 100 nm). After 24 hours all the aggregates appear enclosed in at least a double-membrane, consistent with Wu *et al.*’s observation in Huntington’s disease neurons [29], with some residing within multilamellar structures as well. In 20 DIV neurons, the aggregates remain readily detectable but take on a more variegated morphology (Fig. 1 B.1, B.2), and many appear smaller (Fig. 2 A). Additionally, some 20 DIV neurons appear to no longer be double membrane bound (Fig. 2 A). However, we do still observe aggregates similar to the 1-day aggregates (Fig. 1 C.1, C.2).

**Figure 2.**
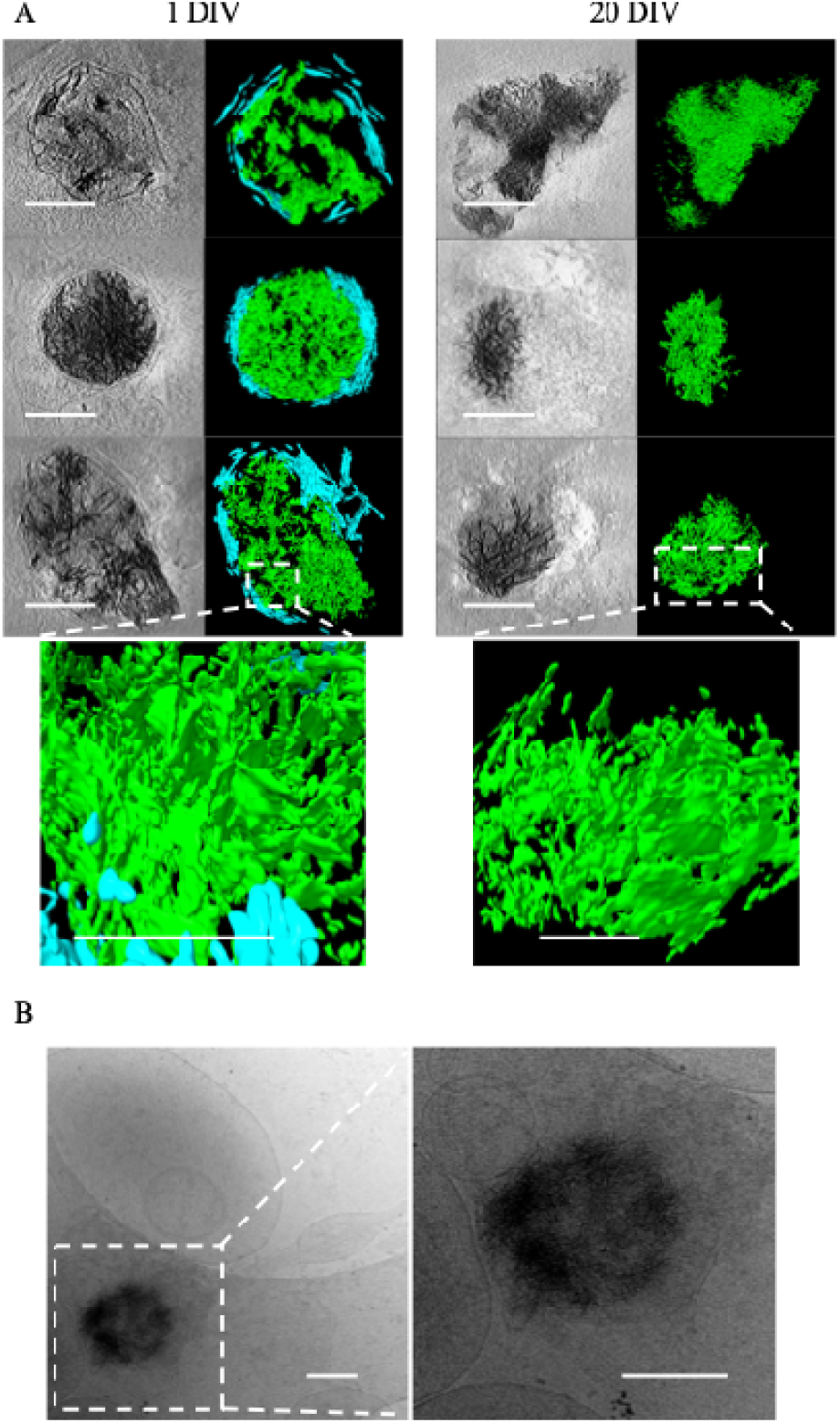
Sheet aggregates show filamentous morphology in 2D and sheet morphology in 3D. **(A)** Cryo-ET tomograms and corresponding segmentations of different kinds of aggregates in 1 DIV rat cortical neurons (left column images) and 20 DIV rat cortical neurons (right column images). Bottom images in each column depict a magnified side view of the sheet morphology in regions delineated by the dashed boxes. In the segmentations, green = aggregates; cyan = membranes. **(B)** Zero tilt micrograph of an aggregate found in a 10 DIV cultured hippocampal neuron. The tilt-series was collected independent of our group for a previously published study [30], demonstrating the aggregates are not unique to our isolation/culturing method. All scale bars = 200 nm except for the bottom magnified side view images in (A), where scale bars = 100 nm.

Wu *et al.* only detected aggregates in their Huntington’s disease model human induced pluripotent stem cells (iPSCs) and Huntington’s disease model primary mouse cortical neurons. They did not observe aggregates in WT control cells [29]. To ensure the aggregates are not specific to our group’s isolation method, we viewed embryonic rat hippocampal neuron tilt-series produced and collected independently and were able to identify aggregates in 10 DIV neurons (Fig 2 B). While aggregates were less common in these tomograms, the images were collected for a publication [30] focused on axons and dendrites, and so cell bodies and extracellular regions were intentionally avoided. Similarly, before our focus shifted to specifically studying aggregates, they appeared only occasionally, as we were primarily focusing on synapses, where they occur infrequently. Since most past Cryo-ET studies of neurons have been focused on the thinnest, easiest to image regions, this may explain why aggregates have not been previously reported in WT neurons.

Establishing the exact chemical identity of the aggregates has required multiple strategies. Wu *et al.* noted they do not have any clear periodic structure in Fourier space or a similar structure to previously known protein aggregates like IZ-synuclein, β-amyloid, or Huntingtin. Additionally, they do not appear morphologically similar to any proteins known to form sheet structures such as clathrin and spectrin. The high electron density of the aggregates, comparable to amorphous calcium-phosphate mitochondrial granules, indicates that a large fraction of the aggregate mass must be a high-Z element. We hypothesized that if Ca^2+^ is a major component of the aggregates, then chelation would cause some change in the aggregates. As a simple test, we chelated 1 DIV neuron media with 10 mM EGTA for 5 minutes prior to vitrification, and found that, despite the short timeframe, the sheet structure of many of the aggregates was ablated as compared to control neurons (Fig. 3 A). All micrographs were collected with the same electron dose and defocus. While the ablated aggregates still retained some amorphous hyper-electron dense material, the fibrous appearance in 2D is greatly reduced to the extent that identification even becomes difficult.

**Figure 3.**
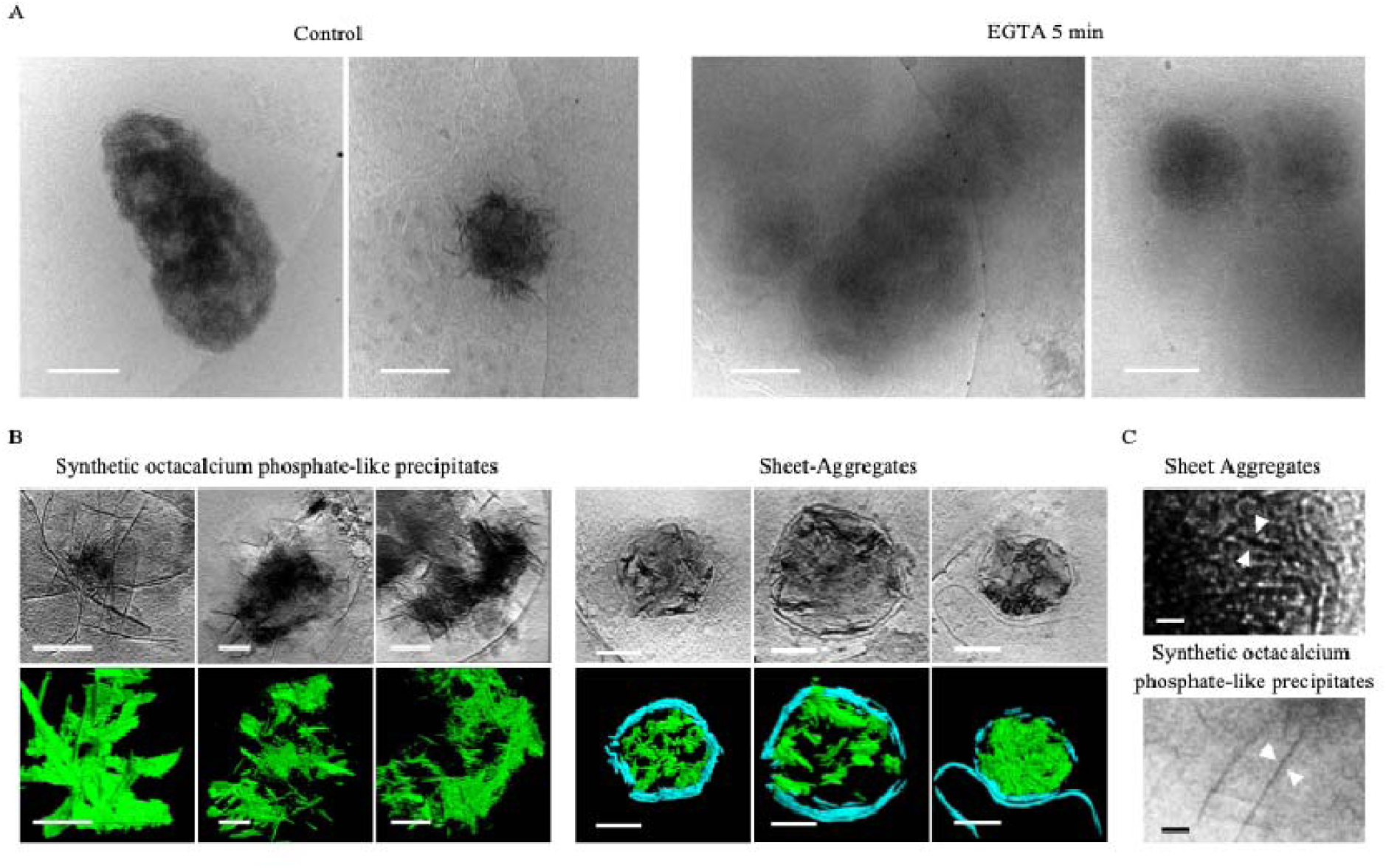
Sheet aggregates have a calcium-dependent structure and appear morphologically similar to OCP-like precipitates. **(A)** Cryo-ET tomograms and corresponding segmentations of 1 DIV cortical neurons incubated in low-magnesium artificial cerebrospinal fluid and either immediately frozen (left images) or incubated with 10mM of EGTA for 5 minutes (right images). **(B)** Cryo-ET tomograms and corresponding segmentations depicting OCP-like precipitates and the aggregates. In segmentations, green = OCP-like precipitate/aggregates; cyan = membranes. **(C)** Magnified view of OCP-like and aggregate micrograph depicting their fibrous appearance in 2D. All scale bars = 200 nm except in (D), scale bars = 10 nm.

ACP is frequently the first CaP phase to precipitate in cellular environments like the mitochondria because, despite HA being favored in the slightly alkaline mitochondrial matrix pH of 7.8-8 (Wilson et al., 2021), ions such as magnesium and citrate inhibit its conversion [16]. However, if the Ca:P ratio is maintained at ∼1.67, the pH >∼7.0, and the inhibitory ion concentrations (e.g. Mg^2+^) permit, HA will form over time. If instead the ratio and pH drop to ∼1.33 and ∼5.5-7.4, OCP will form before HA, and if it continues to drop to 1:1 and <∼5.5, a type of aggregate called brushite will precipitate [15,32]. OCP is striking because it forms ∼1.85 nm thick plates that appear like sheets in 3D [32]. Wu *et al.* found their Huntingtin disease aggregate sheets to be ∼2 nm thick, and we found a similar range for ours, though because the sheets often are not oriented perpendicular to the field of view and Z-resolution is diminished in tomograms, they can appear to be upwards of 7 nm. Based on this morphologic data, the EGTA experiment demonstrating calcium-dependence, and the aggregates’ striking hyper electron-dense resemblance to ACP mitochondrial granules, we hypothesized that these aggregates may represent OCP.

To test this hypothesis, we chemically synthesized OCP aggregates and performed cryo-ET for comparison with the cellular aggregates. Habraken *et al.* previously reported OCP-like structures form when CaCl_2_ is mixed slowly with K_2_HPO_4_ in Tris buffered saline at room temperature and pH 7.4 [32]. In the first ∼20 minutes, dendritic-like pre-nucleation calcium triphosphate ([(Ca(HPO_4_)_3_]^4-^, Ca:P 0.30 – 0.40) complexes aggregate into polymeric strands/nodules (Ca:P 0.30 – 0.40). After ∼20-60 minutes these rearrange into Ca^2+^-deficient amorphous calcium phosphate ([Ca_2_(HPO4)_3_]^2-^ Ca:P ∼0.67) spheres, then Ca^2+^-deficient OCP-like aggregated spheres and ribbons (15-110 minutes, Ca:P ∼1.00), followed by elongated OCP-like plates (>3 hours, Ca:P ∼1.33), and finally HA plates (1 month+, Ca:P ∼1.67) [32]. The term “OCP-like”, rather than just “OCP”, is used because they contain all the characteristic diffraction peaks for OCP except the (100) peak. This is present when 1.85 nm thin crystal plates form; however, Habraken *et al* reported their plates were slightly thinner at ∼1.4 nm. The calcium deficient ACP was reasoned to generate because it was ACP in solution, and it was noted that when dried, it had the characteristic Ca:P of ∼1.5. This point emphasizes why cryogenic investigation of these aggregates is important, because how the sample is prepared can affect the analysis of calcium-dependent structures. We followed their procedure for 3 hours to enrich for OCP-like ribbons and plates. We then vitrified the samples, collected tomograms of identified precipitates, segmented the structures, and compared them to the aggregates.

Both the aggregates and OCP form fibrous strands in 2D, which are clearly sheets when segmented in 3D (Fig. 3 B). Though their fiber thicknesses appear similar, close inspection does reveal visual differences (Fig. 3 C). Some of the aggregates appear mottled, and generally the aggregate clumps are smaller. This is likely due to the fact that when formed in neurons, they are membrane-bound, which confines the size and direction the sheets can grow. Additionally, in the aggregates’ environment competing ions like Mg^2+^, carbonates, and other phosphates can bind, thereby affecting the sheet thickness by changing how planes stack.

To confirm they have an OCP-like structure, we used LDSAED to compare the diffraction pattern of aggregates from 1 DIV neurons to the lab synthesized OCP-like elongated plates. Representative diffraction images for both samples show broad, diffuse rings consistent with poorly ordered/non-crystalline material (Fig. 4A). Habraken *et al.* previously reported OCP-like ribbons and elongated plates have a broad peak between ∼2.6-3.3 Å that changes to a sharp peak at 2.8 Å as it converts to HA [32]. To investigate whether this is present in our data, we collected images with an aggregate centered in the image (“full”), where only a small amount of aggregate was visible (“partial”) and a nearby control background region. We then divided the radially averaged intensity distribution for the “full” and “partial” images by each sample’s corresponding background to dampen non-specific peaks such as those at ∼3.71 □ and ∼2.15 □ due to vitreous ice [33]. We found a broad peak ranging between ∼2.5-3.2 Å present in the “full” image that overlaps well with our OCP-like elongated plate control (Fig. 4A, 4B) and to the previously reported 2.6-3.3 Å peak observed. This supports the sheet aggregates being HA precursor OCP-like plates. We next investigated where these aggregates specifically nucleate in neurons.

**Figure 4.**
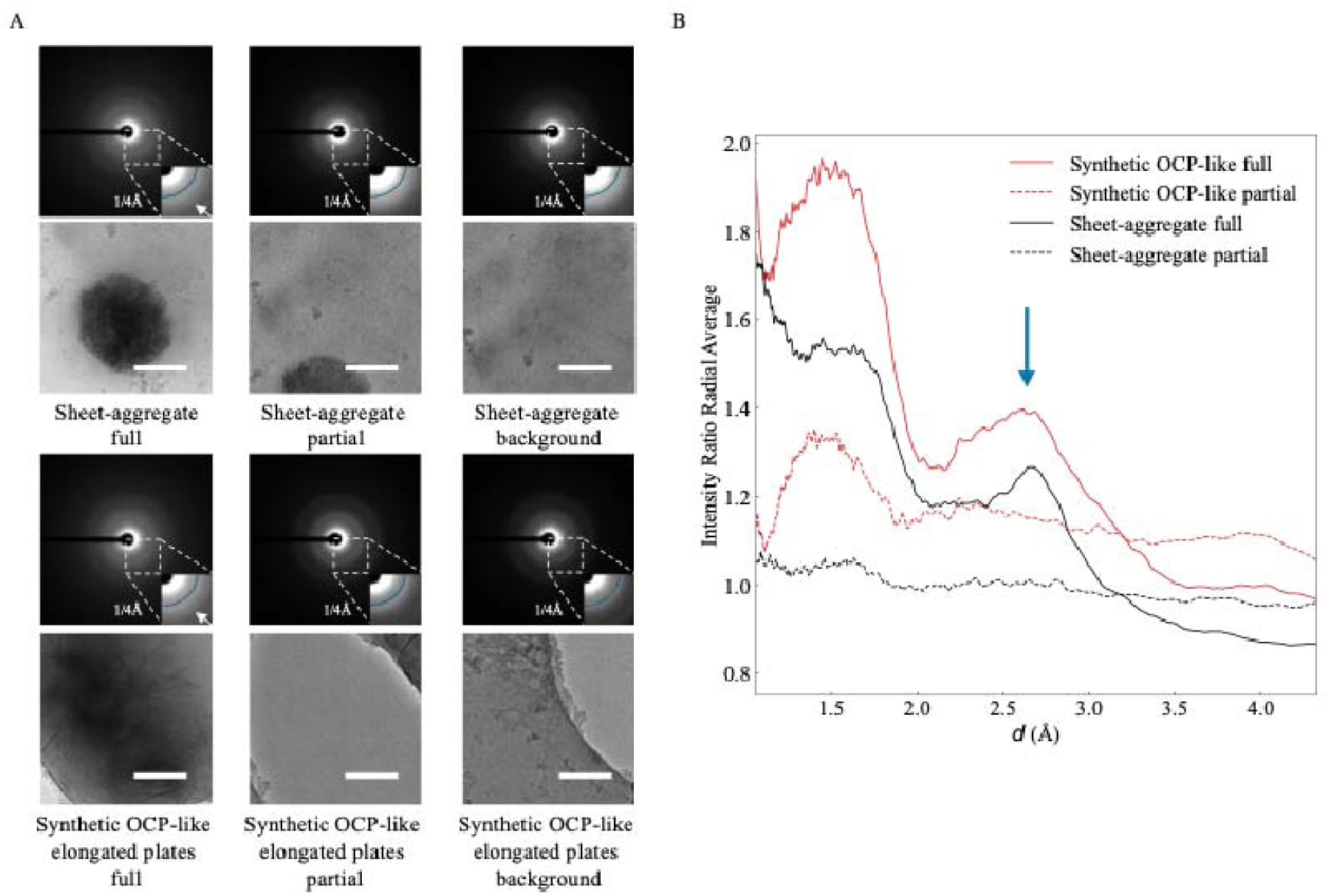
Sheet aggregates have a similar diffraction signature to OCP-like elongated plate HA precursors. **(A)** LDSAED of a sheet aggregate from 1 DIV rat cortical neurons and synthesized OCP-like elongated plates. Top image of the pair is the electron diffraction micrograph depicting a maximum spatial frequency of 1/1.0 □^-1^in the full image and 1/2.4 □^-1^in the inset. Reference radial curve in the inset image corresponds to 1/4 □^-1^. The white arrow is emphasizing the broad band between ∼1/2.6 □^-1^ – 1/3.3 □^-1^ observed previously in OCP-like elongated plates [32]. Scale bars = 200 nm. **(B)** Ratio of radially averaged full or partial electron diffraction image intensities from (A) over each group’s respective background image. The x-axis has been converted to d spacing (1/q) and the arrow indicates the broad ∼2.6-3.3 □ peak.

Given that many of the aggregates are found extracellularly 1 day following isolation, we hypothesized they are a deleterious CaP precipitate that the cells clear after ionic disequilibrium occurs. In line with this thinking, Wu *et al.* conjectured previously that the aggregates could be derived from damaged mitochondria undergoing mitophagy [29]. To test if the aggregates are associated with mitochondria, we stained for a protein member of the translocase of the outer mitochondrial membrane complex (TOM) called TOM20. We achieved this by fixing cells in paraformaldehyde and performing immunofluorescent staining prior to freezing (Fig. 5 A-F). Upon visualization, we found many aggregates were associated with the TOM20 stain (Fig. 5 C-E) but not all (Fig. 5 F). Intriguingly, whether the aggregates were found intracellularly or extracellularly was a significant predictor of its TOM20 status. We found ∼92% of intracellular aggregates identified stained positive for TOM20 versus only ∼61% of extracellular aggregates (Fig. 5 G). Additionally, we found nascent aggregates forming within visually identified mitochondria (Fig. 5 E), providing strong evidence for mitochondrial origin, though we cannot exclude the possibility of other additional sources. Next, we immunostained neuron cultures for the electron transport chain protein COX4 (Fig. 5 H-J) and found that mitochondria both with and without calcium-phosphate granules stain positive (Fig. 5 I and J). However, the aggregates do not appear associated with COX4 (Fig. 5 J). The findings that most intracellular aggregates stain positive for TOM20 and nascent sheets can be found in mitochondria support the hypothesis that at least some aggregates are of mitochondrial origin. The result that they do not stain positive for respiratory proteins like COX4 suggests these aggregates reside within mitochondria that are no longer functional, or they are within vesicles derived from the mitochondrial membrane.

**Figure 5.**
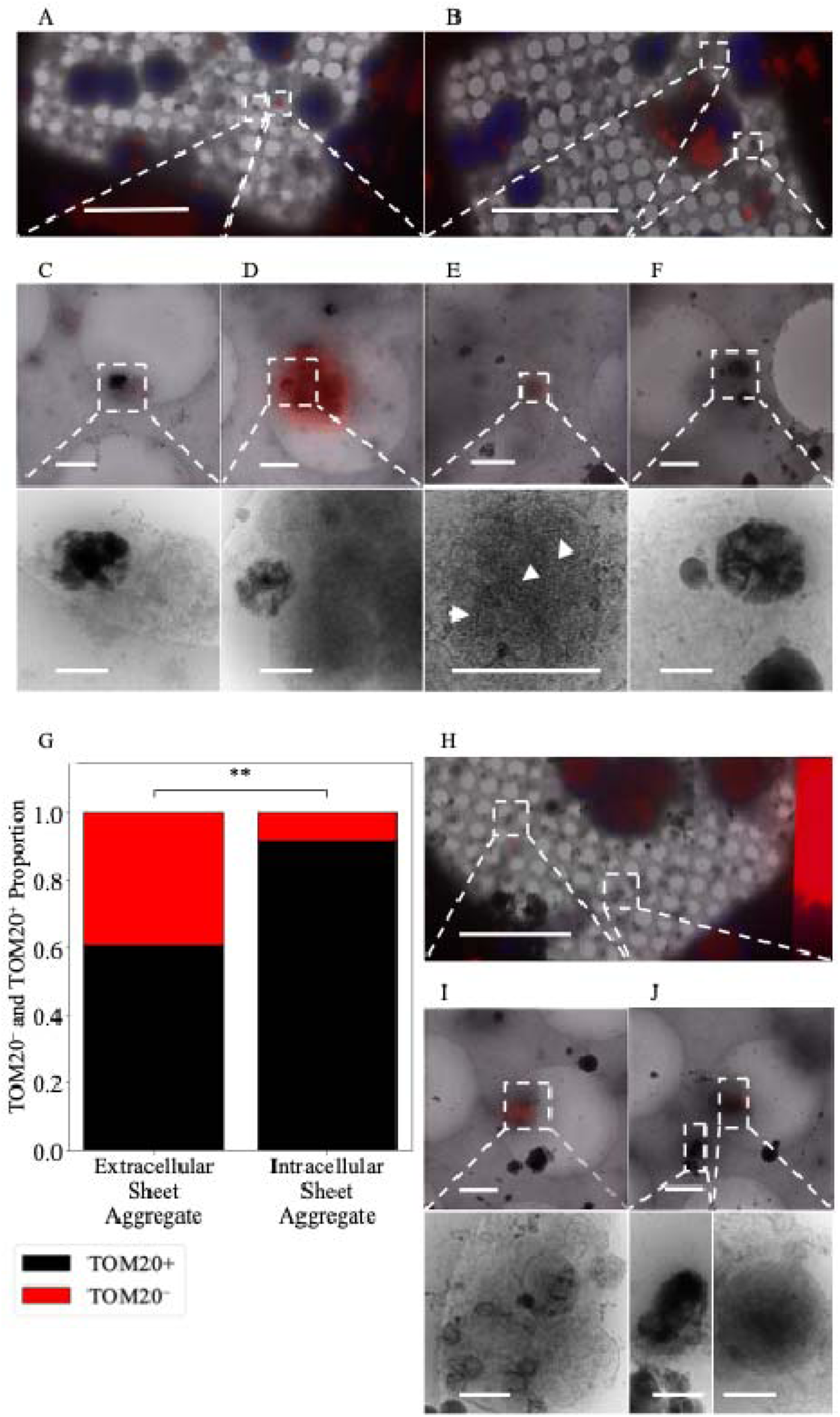
Sheet aggregates are well associated with TOM20^+^ mitochondria but poorly associated with COX4^+^ mitochondria. **(A)** and (**B)** 700× micrographs of PFA fixed vitrified 1 DIV rat cortical neurons overlaid with TOM20 (red) and Hoechst (blue) fluorescent stains. (**C)** 5,300× micrograph of extracellular aggregates with a weak TOM20 stain (top) and corresponding 45,000× image (bottom). **(D)** 5,300× micrograph of aggregates with a strong TOM20 stain near TOM20^+^ mitochondria (top) and corresponding 45,000× image (bottom). **(E)** 5,300× micrograph of aggregates forming within TOM20^+^ mitochondria (top) and corresponding 45,000× image (bottom), white arrowheads depict nascent sheets forming. **(F)** 5,300× micrograph of extracellular aggregates with no TOM20 stain (top) and corresponding 45,000× image (bottom). **(G)** Plot depicting the proportion of aggregates that are extracellular TOM20^+^ (n=28, 61%) or TOM20^-^ (n=18, 39%) versus the proportion that are intracellular TOM20^+^ (n=33, 92%) or TOM20^-^ (n=3, 8%). A two-sided Fisher’s exact test was used to determine significance; odds ratio = 0.141, p-value=0.001883, 95% CI = (0.05 - 0.56). ** = p-value ≤ 0.01. **(H)** 700× micrographs of PFA fixed vitrified 1 DIV rat cortical neurons overlaid with COX4 (red) and Hoechst (blue) fluorescent stain. **(I)** 5,300× micrograph of COX4^+^ extracellular mitochondria with calcium phosphate granules (top) and corresponding 45,000× image (bottom). **(J)** 5,300× micrograph of extracellular COX-negative aggregates and COX4^+^ mitochondria (top) and corresponding 45,000× image of aggregates (bottom left) and mitochondria (bottom right). Dashed boxes and lines depict areas of interest that are gradually increased in magnification. Scale bars for 700× micrograph = 25 µm; 5,300× = 1 µm; and 45,000× = 200 nm.

As previously discussed, Wu *et al.* hypothesized the aggregates may be derived from mitochondria undergoing mitophagy [29]. To test for mitophagy involvement, we stained for lysosomes using Lysotracker^TM^ as well as the ATG8 family protein LC3, which is a ubiquitous marker for autophagy/mitophagy during protein cargo recruitment and autophagosome-lysosome fusion [34]. We did not detect the aggregates spatially associated with lysosomes or LC3 (Fig. 6 A-F). Because the aggregates do not appear to be within lysosomes or LC3^+^ autophagosomes, given many are found extracellularly and migrasomes have been shown to remove damaged mitochondria [22], we hypothesize they may be extruded through mitocytosis.

**Figure 6.**
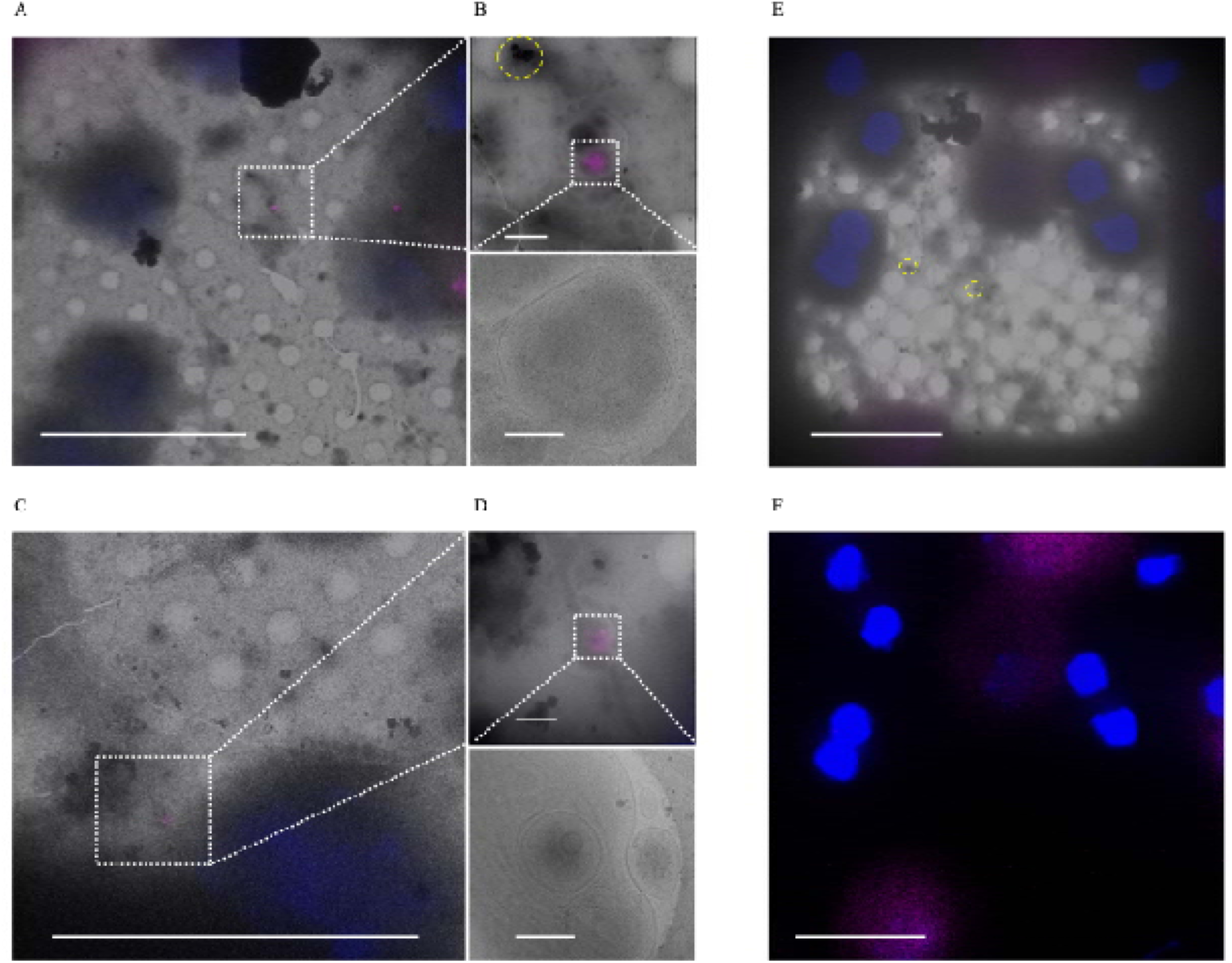
Sheet aggregates are not associated with lysosomes or LC3^+^ autophagosomes. **(A)** and **(C)** 700× cryo-EM micrographs of PFA fixed rat cortical neurons overlaid with Lysotracker^TM^ (purple) and Hoechst (blue) fluorescent stains. **(B)** and **(D)** 5,300× cryo-EM micrographs of lysosomes overlaid with Lysotracker^TM^ and Hoechst stains (top images) and their corresponding 45,000× micrographs (bottom images). **(E)** 700× cryo-EM micrographs of PFA fixed rat cortical neurons overlaid with LC3 (purple) and Hoechst (blue) fluorescent stains. Dashed boxes and lines depict areas of interest that are gradually increased in magnification. **(F)** Same region as **(E)** except depicting the fluorescent signal alone without the micrograph overlay to emphasize the lack of LC3 puncta in a 700× micrograph with aggregates. Yellow dashed circles are outlining aggregates. Scale bars for 700× micrograph = 20 µm; 5,300× = 1 µm; and 45,000× micrograph = 200 nm.

Recently, the metalloprotease CPQ was identified as a highly specific marker for migrasomes that differentiates them from other extracellular vesicles like exosomes [35]. We stained for CPQ in our 1 DIV cortical neurons and identified aggregates inside CPQ^+^ extracellular vesicles (Fig. 7 E-H). We also found CPQ^+^ vesicles containing mitochondria (Fig. 7 G), which supports the previous finding that migrasomes undergo mitocytosis [22]. CPQ was not associated with all extracellular aggregates (Fig. 7 I and J). However, in these cases the aggregates no longer appeared to be inside any larger migrasome-appearing vesicle, implying they may be byproducts of migrasomes that have released their contents [24].

**Figure 7.**
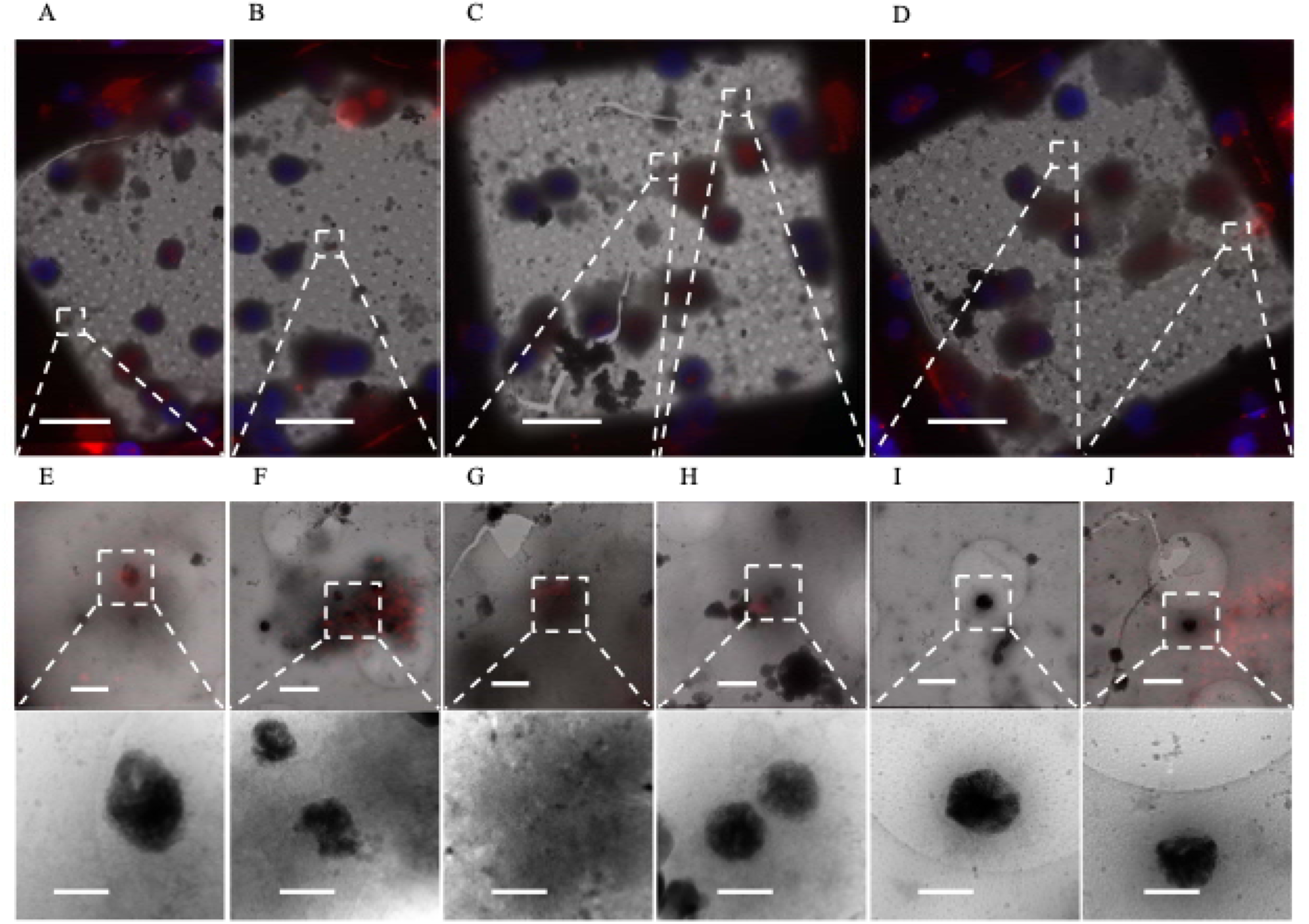
Sheet aggregates are found in CPQ^+^ migrasomes. **(A-D)** 700× cryo-EM micrographs of PFA fixed vitrified 1 DIV rat cortical neurons overlaid with CPQ (red) and Hoechst (blue) fluorescent stains. **(E)**, **(F)**, and **(H)** 5,300× cryo-EM micrographs depicting aggregates inside CPQ^+^ migrasomes (top) and their corresponding 45,000× images (bottom). **(G)** 5,300× cryo-EM micrograph depicting mitochondria inside CPQ^+^ migrasomes (top) and its corresponding 45,000× images (bottom). **(I)** and **(J)** 5,300× cryo-EM micrographs depicting extracellular aggregates not associated with CPQ^+^ migrasomes (top) and their corresponding 45,000× images (bottom). Dashed boxes and lines depict areas of interest that are gradually increased in magnification. Scale bars for 700× micrograph = 20 µm; 5,300× = 1 µm; and 45,000× = 200 nm.

Knowing the aggregates are likely CaP precipitates that form in mitochondria and given that shocks of high calcium can cause precipitation events, we investigated whether incubation in media with a lower calcium concentration would decrease their size. We incubated cells in Neurobasal (NB) or F-12 media +/- Fetal Bovine Serum (FBS) after standard isolation since F-12 has a lower calcium (0.3 mM vs 1.8 mM), glucose (10 mM vs 25 mM), and glutamine (1 mM vs 2 mM) concentration. We also tested the effect that time in culture has on aggregate formation. To investigate their size, we collected >90 2D micrographs of aggregates from all conditions and measured their area. We found that the aggregates in neurons from the 1 DIV F-12 + 10% FBS media and 20 DIV NB media conditions appear to be significantly smaller than the other groups (Fig. 8 A). Interestingly, the decreased area in the 1 DIV F-12 + 10% FBS and 20 DIV NB conditions were different in character from each other. Aggregates from the F-12 condition neuron cultures appeared within membrane boundaries of similar size to the control conditions; however, the aggregates themselves were less dense, leading to a smaller area value (Fig. 8 B). Conversely, aggregates from 20 DIV control neurons appeared similarly dense but overall smaller in size than those found in the 1 DIV control neurons (Fig. 8 B). These data indicate that neuronal media affects the size of the aggregates and longer time in culture increases the number of small aggregates.

**Figure 8.**
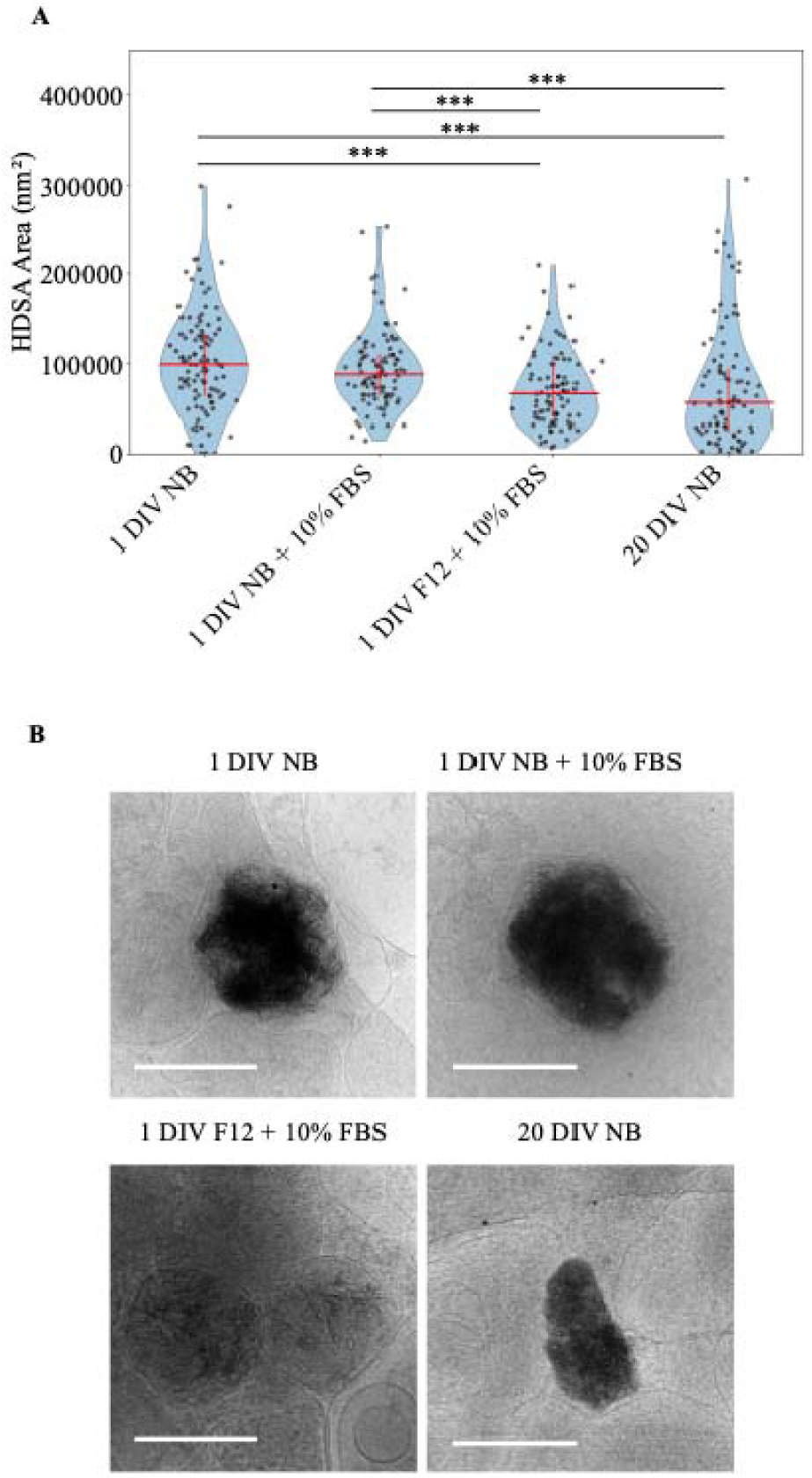
Sheet aggregate area decreases with F-12 media and time in culture. **(A)** Graph depicting area of aggregates in nm^2^ across different conditions. For each condition grids from 3 different cultures were imaged. 1 DIV Neurobasal (NB) media, n = 107; 1 DIV; Neurobasal (NB) + 10% FBS n = 100; 1 DIV F-12 + 10% FBS n = 94; 20 DIV NB n = 94; Median and interquartile range depicted in red. Wilcoxon rank-sum test followed by Benjamini-Hochberg correction for multiple comparison was done. *** = p <0.001, ** = p < 0.01, * = p < 0.05. **(B)** Representatives of median sized aggregates: 1 DIV NB median = 99,318 nm^2^; 1 DIV NB+ 10% FBS, median = 88,134 nm^2^; 1 DIV F-12 + 10% FBS, median = 66,782 nm^2^; 20 DIV NB, median = 56,475 nm^2^. All scale bars = 200 nm.

## Discussion

We have observed the presence of hyper-electron-dense sheet aggregates in both 1 and 20 DIV embryonic rat cortical neurons. They appear both intra and extracellularly, as well as within both neurites and cell bodies. Calcium chelation with EGTA ablates the sheet morphology, but retains high contrast, strongly implicating Ca^2+^ as a critical component of the complex. Electron diffraction analysis further supports the identity of the aggregates as OCP-like plates via a broad band in the ∼2.5-3.2 Å range matching lab-grown CaP aggregates. Together these experiments provide strong evidence that these aggregates are primarily composed of CaP in an OCP-like state.

To assess the cellular origin of the aggregates, we first observed that in both 1 DIV and 20 DIV primary cortical neurons they appear intracellularly, extracellularly, in neurites, and cell bodies, and retain a mitochondria-like appearance and size. After 1 day all the aggregates appeared to be at least double membrane bound, however some aggregates in the 20 DIV group do not show a discernable encapsulating membrane. We stained 1 DIV neurons for the outer membrane mitochondrial protein TOM20 and found many of the aggregates associated with this protein. However, these aggregates do not appear significantly associated with the electron transport chain protein COX4, indicating they may be non-energetically active/damaged mitochondria or mitochondria-derived vesicles. They also do not appear associated with the ATG8 family LC3 protein involved in autophagosome recruitment or lysosomes. However, they are associated with CPQ^+^ extracellular vesicles called migrasomes, which may serve as one of the cell’s mechanisms for extracellular export. Culturing neurons for 1 day in F-12 media + 10% BSA, which contains a decreased Ca^2+^ concentration, or for 20 days in NB media causes the aggregates to decrease in size relative to those in 1 DIV neurons cultured in NB. These findings support the hypothesis that following the metabolically stressful isolation procedure, primary cultured rat cortical neurons form mitochondria-derived OCP-like precipitates.

Formation of these aggregates appears to vary significantly with culturing conditions and buffer. While we can only speculate, this does offer a plausible explanation for why Wu *et al.* detected similar aggregates in human and mouse Huntington’s disease cortical neurons but not their corresponding WT control neurons [29]. Human iPSC neural progenitors, derived from fibroblasts, do not undergo the same challenging isolation procedure. It is less clear why they were not observed in their mouse neuronal cultures, but some differences in our culture conditions do exist. Their embryonic mouse neurons were incubated in Neurobasal + 10% FBS and GlutaMAX^TM^ for a day before switching to serum free media for 13 more days before cryopreservation. Our normal culture protocol uses 2mM glutamine without any serum incubation. While we found 10% FBS after 1 day with Neurobasal does not significantly decrease the aggregate size, we did not follow those cultures for 13 days. Given our findings that longer times in culture and post-isolation media with lower Ca^2+^ concentration cause the aggregates to decrease in size, it is plausible that anything that changes the calcium, phosphate, pH, or inhibitory ion concentration of the media during or following isolation will impact aggregate solubility and structure. Overall, our findings suggest that while Huntington’s disease likely causes an increased presence of these aggregates, they are not limited to this disease state as previously proposed.

It is well established that ACP granules can precipitate within mitochondria [12,36]. However, the presence of OCP-like structures in mitochondria remains more controversial, as some have argued that mitochondrial ionic conditions are not conducive to OCP formation [36]. Despite this, the striking morphological and structural similarities between the intracellular aggregates we observed and the laboratory synthesized OCP-like material presents a compelling case. While definitive elemental and structural characterization, such as cryo-energy dispersive X-ray spectroscopy (cryo-EDX) or electron energy loss spectroscopy (cryo-EELS), could further strengthen this interpretation, access to these techniques for cryo-preserved specimens is currently limited to a small number of international sites and room temperature protocols are highly perturbative.

Our findings fit well into previous studies that investigated *in vivo* CaP formation in the nervous system. Hydroxyapatite was reported in the mitochondria of mouse neurons from fixed cortical tissue following excitotoxic shock [18], and microcalcification in Alzheimer’s disease patient neurons was associated with phosphorylated Tau presence [37]. These demonstrate hydroxyapatite can form *in vivo* under pathologic conditions, but these and the vast majority of past studies were done using structure-altering fixatives and stains to view tissue samples at room temperature. In recent years however, cryo-FIB/SEM opens the possibility of investigating brain tissue and slice cultures in a near-native environment, and we predict as this method is increasingly used these OCP-like aggregates will be more widely observed.

Mitochondria have been implicated as a source of CaP during bone formation [13,14]. Extending this concept to neurons, our findings reveal that large OCP-like aggregates, substantially larger than the typical mitochondrial CaP granules, can form within mitochondria and are capable of being exported into the extracellular space. These results suggest a potential mechanism linking mitochondrial dysfunction to brain calcification disorders. Although our observations are in neurons, they bear striking parallels to CaP dynamics in osteoblasts during bone mineralization. This raises the intriguing possibility that brain calcification may arise, at least in part, from aberrant activation of pathways normally involved in bone growth. More broadly, our findings suggest that metabolically stressed neurons can act as a nucleation site for CaP accumulation, providing a mechanistic explanation for its extracellular deposition in neuropathologies such as the β-amyloid plaques of Alzheimer’s disease [38]. These insights further support growing interest in targeting mitochondrial dysfunction as a therapeutic strategy in neurological diseases.

## Methods

### Neuron Culture and Media Experiments

Neuron extraction was performed as previously described [25,26]. 1 day prior to isolation 2-4 cryoEM grids were placed in 35 mm Matsunami glass bottom culture dishes (Avantor Sciences) dishes and UV-sterilized in the cell culture hood for 30 minutes. Quantifoil R 2/2 Au 200 mesh and R 3.5/1 200 mesh Au cryoEM grids were used. Then, 0.1 mg/mL poly-D lysine (Thermo Fisher) was added to the dish and incubated overnight at 37° C 5% CO_2_. The next day, poly-D lysine was removed and the plates were washed 3× with phosphate buffered saline (PBS). After isolation, neurons were cultured at 37° C and 5% CO_2_ in Neurobasal media (Thermo Fisher) supplemented with 50X B27 (Thermo Fisher), 2mM L-glutamine (Thermo Fisher) and 10,000 U/mL Penicillin-Streptomycin (Thermo Fisher).

To test the effect of fetal bovine serum (FBS, Thermo Fisher) and F-12 media (Thermo Fisher), following isolation cells were either incubated with NB alone, NB + 10% FBS, or in F-12 media + 10% FBS. Cells incubated for 24 hours did not have their media exchanged prior to freezing. Cells incubated for 20 days underwent a full media exchange 24 hours after plating, and then every 3-4 days 1/3^rd^ of the media was removed and replaced with fresh media. To test EGTA chelation, 23 hours after plating cells were incubated in low magnesium 0.2 µm-filtered 1.5X artificial cerebrospinal fluid (aCSF) pH 7.4 for 1 hour, containing 186 mM NaCl, 4.5 mM KCl, 1.0 MgCl_2_, 15 mM D-glucose, 15 mM HEPES, 4.5 mM CaCl_2_ (all reagents from Sigma-Aldrich). 10mM EGTA was then added for 5 minutes and neutralized by removing the media, washing 3X with aCSF and replacing with fresh media. Neurons were then frozen within 10 minutes of stimulation.

### Sample Vitrification

Neurons were frozen in liquid ethane cooled by liquid nitrogen using the Leica EM GP plunge freezer. Owing to transfer, room temperature incubation prior to freezing is unavoidable but was limited to less than 10 minutes. The chamber was set to 60% humidity, and 1 DIV neuron grids were back blotted for 8 seconds. 20 DIV neuron grids were back blotted for 15 seconds. 1.5 µL of 1:1 fresh Neurobasal media:cultured Neurobasal media was added to the front of the grid prior to blotting, except for EGTA stimulation where just fresh aCSF was used. Octacalcium phosphate precipitates were vitrified on R 1.2/1.3 Cu 200 mesh Quantifoil grids. The grids were glow discharged first using the PELCO easiGlow^TM^ with 15 mAmps for 30 seconds, a 15 second hold time, and 0.4 mBar pressure. The grids were similarly frozen with the Leica EM GP plunge freezer except 3 µL of solution was added to the front of the grid and back blotted for 8 seconds. All grids were clipped with autogrid rings and C-clips (Thermo Scientific and Ted Pella) and stored in liquid nitrogen until imaging as described below.

### Cryo-Electron Microscopy and Tomography Collection

Frozen clipped neuron grids were imaged using a 200kV Glacios^TM^ (Thermo Scientific), Falcon 4i direct electron detector (Thermo Scientific), and Tomography 5 software (Thermo Scientific). For cryo-ET, tilt series were acquired with an exposure of ∼1.2-1.8 e^-^/□^2^/tilt and total dose of ∼61-91 e^-^/□^2^/tilt series. We used a dose symmetric collection scheme ranging between -50° and +50° with 2° tilt step between -3 µm to -7 µm defocus and at 3.12 □/pixel or 1.24 □/pixel. EMAN2 was used to reconstruct tomograms using the pipeline described previously [39]. Segmentations were created using EMAN2’s semi-automatic convolutional neural network method described previously [39]. Segmentations were manually cleaned in ChimeraX [40] using their Hide Dust and Map Eraser plugins to remove false positives generated by the neural network. Each segmentation and corresponding tomogram for display were then Gaussian filtered with a width of 20 using ChimeraX’s Map Filter plugin. For the EGTA chelation experiment, images were also collected on the Glacios^TM^ with 5 e^-^/ □^2^, -5 µm defocus, and 3.21 □/pixel for each image. The experiment was repeated twice and grids from two different cultures for each condition (EGTA chelation and aCSF control) were imaged.

### Immunofluorescent Staining

Neurons were grown as described above for 24 hours, however prior to vitrification cells were fixed and stained. When staining for lysosomes, prior to fixation we incubated with LysoTracker^TM^ (Thermo Fisher, L12492) at 50 nM for 30 minutes at 37° C and 5% CO_2_. To fix cells, we removed the Neurobasal media and washed 1× with PBS containing calcium and magnesium (Thermo Fisher). We then fixed the cells in 4% formaldehyde (Sigma-Aldrich) and diluted in PBS for 15 minutes at room temperature. Cells were then washed 3× with PBS and permeabilized for 10 minutes at room temperature using PBS containing 0.25% Triton X-100 (Sigma-Aldrich) and washed again 3× with PBS. Next, cells were blocked for 1 hour at room temperature with 5% bovine serum albumin (BSA, Sigma-Aldrich) in PBS and stained with the primary antibody overnight diluted in the 5% BSA blocking solution at 4°C (antibody information below). The next day, cells were washed 3× with 0.1% Tween-20 (Sigma-Aldrich) in PBS with 5-minute incubation between each wash, and then the secondary antibody was added, diluted in the blocking solution, and incubated at room temperature for 1 hour away from light. The cells were then washed 3× with PBS + 0.1% Tween-20, again with 5-minute incubations between washes, then washed 1× with regular PBS, and then vitrified as described above.

We used the following primary and secondary antibodies and dilutions. Primary antibodies: Rabbit TOMM20 [EPR15581-54] 1:250 (Abcam, ab186735); Rabbit COX IV 1:200 (Abcam, ab153709); Mouse LC3B [G-9] 1:100 (Santa Cruz Biotechnology, sc-376404); Rabbit PBCP/CPQ 1:100 (Proteintech, 16601-1-AP). Secondary antibody: Goat anti-rabbit IgG Alexa Fluor^TM^ 565 nm 1:500 (Thermo Fisher, A11011); Donkey anti-mouse IgG Alexa Fluor^TM^ 647 nm 1:500 (Thermo Fisher, A32787). To stain nucleic acids, we used Hoechst 33342 at 1 µg/mL (Thermo Fisher, 62249).

### Immunofluorescent Imaging

Fixed and frozen neurons were imaged using the Aquilos 2 Cryo-Focused Ion Beam/Scanning Electron Microscope (cryo-FIB/SEM) equipped with the Integrative Fluorescent Light Module (iFLM) (Thermo Scientific). For this experiment only the SEM and iFLM were used. Grids were loaded into the microscope using the 35° shuttle and mapped with the MAPS 3.24 software (Thermo Scientific) using 195× magnification, 2kV voltage and 13pA current. When an ideal region was identified on the SEM map, the grid was imaged with the iFLM using the iFLM 1.3 software. The 385 nm channel was used to collect the Hoechst stain with an intensity of 10% and exposure time of 50 ms. The 565 nm channel was used for the Alexa Fluor^TM^ 565 nm secondary antibody with 20% intensity and 50 ms exposure time. A reflection image for alignment was also collected using 1% intensity and 0.1 ms exposure time. A z stack of images was taken using a slice distance of 0.7 µm and the stack was summed into a maximum intensity image used for identification. 1-3 regions were collected per grid depending on how many sufficient regions were identified. Regions were then overlaid with the SEM map using the program’s semi-automatic alignment method. Then the grid was transferred to the Glacios^TM^ for high-resolution imaging. An overview image at a magnification of 28.3 µm/pixel was taken and overlaid onto the SEM/fluorescent image in the MAPs program using the Hoechst nuclear stain as anchor points. Fluorescent puncta of interest were then identified and imaged in the Glacios^TM^ at a pixel size of 3.2 □/pixel with 10 e^-^ total dose and -10 µm defocus, and a search magnification of 13.5 □/pixel and -85 µm defocus. When quantifying the proportion of intracellular vs extracellular aggregates that are TOM20^+^, we used a two-sided Fisher’s exact test to determine significance.

### Aggregate Area Calculation

To calculate aggregate areas, neurons were grown in each condition and vitrified as described under the “Neuron culture and media experiments” and “Sample vitrification” methods sections. Using the Glacios^TM^ we collected ∼30 micrographs of aggregates from 3 different cultures of each condition at a defocus of -10 µm, 3.21 □/pixel, and 10 e^-^/image. Segmentation maps were manually generated using EMAN2 over each aggregate area [41]. Using custom python scripts each mask was multiplied by the original image so only the aggregate pixel area remained. Each aggregate’s area within the image was then binary thresholded based on the equation: (*maximum pixel value* – *minimum pixel value*) – *pixel standard deviation ×* 1.5 for that specific area. E.g., if 2 aggregates were in the same image, a different threshold value for each aggregate was determined based off its pixel distribution. This was chosen empirically by visualizing all the images with different threshold levels and determining what maintained the most aggregate pixels over background. The area was determined using the Python skimage.measure function regionprops, converted to nm^2^, and graphed. Statistical significance was calculated using the Wilcoxon rank-sum test and p-values were corrected for multiple comparison using Benjamini-Hochberg correction.

### Octacalcium Phosphate Precipitation

To precipitate octacalcium phosphate (OCP) we followed the procedure previously reported by Habraken *et al*. [32]. Briefly, a 30 mL working buffer of 50mM Trizma^®^ base (Sigma) and 150 mM NaCl (Sigma) was made with 18.2 MΩ·cm ultrapure water and adjusted to pH 7.4 with 0.1 M NaOH (Sigma) and 0.1M HCl (Sigma). Next, 25 mL 14 mM K_2_HPO_4_ and 5 mL 25 mM CaCl_2_・H_2_O solutions were created in the Tris salt buffer and pH adjusted to 7.4 with 0.1 M NaOH. Then, the CaCl_2_ solution was added to the 25mL KH_2_PO_4_ solution at a rate of 10 µL/min for 3 hours at 21° C under constant 200 RPM stirring. The pH was titrated with 0.1 M NaOH and 0.1 M HCl during the process to maintain a pH of 7.4. After 3 hours, the solution was added to grids, vitrified, and imaged as described in the “Sample vitrification” and “Cryo-electron microscopy and tomography collection” methods sections.

### Low Dose Selected Area Electron Diffraction

Cryo-selected area electron diffraction (SAED) was performed using a 200kV Glacios^TM^ (Thermo Scientific) equipped with a Ceta-D camera (Thermo Scientific). We used a camera distance of 670 mm (Nyquist spatial frequency = 1.75 □^-1^), a SA aperture size of 10 µm, and a defocus of -0.5 µm. Non-diffraction images of areas were also collected using a pixel size of 1.62 □/pixel and a defocus of -2 µm. 1 DIV neuron aggregates and 3-hour OCP-like elongated plates were imaged in an incremental manner beginning directly over the object and gradually moving off to capture how the pattern changes as the target aggregate or OCP is no longer centered. Diffraction patterns were analyzed using EMAN2 [42] and custom python scripts. Specifically, diffraction images were centered, radially averaged, and normalized on the spatial frequency (q) bands 0.094 - 0.113 □^-1^. To correct the background an adjacent image to the target (e.g. aggregate or OCP-like elongated plate) was collected, and the target radial distribution was divided by this distribution.

## Abbreviations

ACP: amorphous calcium phosphate
ATG8: autophagy-related protein 8 (family)
BSA: bovine serum albumin
CaP: calcium phosphate
COX4: cytochrome c oxidase subunit 4
CPQ: carboxypeptidase Q
cryo-CLEM: cryo-correlative light and electron microscopy
cryo-EM: cryo-electron microscopy
cryo-ET: cryo-electron tomography
DIV: *days in vitro*
E18: embryonic day 18
EGTA: ethylene glycol-bis(β-aminoethyl ether)-N,N,N′,N′-tetraacetic acid
F-12: Ham’s F-12 nutrient mixture
F-actin: filamentous actin
FBS: fetal bovine serum
FIB/SEM: focused ion beam / scanning electron microscopy
HA: hydroxyapatite
HBSS: Hanks’ balanced salt solution
iPSC: induced pluripotent stem cell
LC3: microtubule-associated protein 1 light chain 3 (ATG8 family member)
LDSAED: low-dose selected-area electron diffraction
mPTP: mitochondrial permeability transition pore
NB: Neurobasal (medium)
OCP: octacalcium phosphate
PFA: paraformaldehyde
RPM: Revolutions per minute
TOM20: translocase of the outer mitochondrial membrane 20-kDa subunit

## Ethics declarations

## Ethics approval

E18 Long-Evans rats were used for cortical neuron cultures and experiments were approved by the Institutional Animal Care and Use Committees at Baylor College of Medicine (*animal protocol number: AN-4365)*.

## Consent for Publication

Not applicable.

## Data Availability

The datasets generated and analyzed during the current study are available in the Electron Microscopy Data Bank (EMDB) and Electron Microscopy Public Image Archive (EMPIAR). Specifically, representative 1 DIV aggregate tomograms are located on EMDB at accession number: EMD-71853, EMD-71859, EMD-71860, EMD-71861; 1 DIV mitochondria with ACP granules at: EMD-71852; and 20 DIV aggregate tomograms at: EMD-71850, EMD-71851, EMD-71862, EMD-71863, EMD-71864. All files for this project, e.g. the raw data for the cryo-ET, cryo-CLEM, and LDSAED experiments as well as Python scripts used for analysis, can be found on EMPIAR at accession number: EMPIAR-12905.

## Acknowledgements

We thank Dr. Zhao Wang, Isaac Forrester, Robyn Tebbetts, and Juwayriyah Sana Qureshi from the Baylor College of Medicine Cryo-EM core for their electron microscope training and advice. We thank Andrew Yang and Dr. Theodore Wensel for experimental advice and discussions.

## Author Information

## Contributions

ED Anderson conceived the project, collected the data, and wrote the manuscript. CA Cronkite, CP Abella, JG Duman, and AN Simmonds isolated the rat cortical neurons and provided tissue culture and experimental advice. PR Baldwin, SJ Ludtke, and ED Anderson analyzed the data. MN Waxham provided independently collected rat hippocampal neuron tomograms containing the aggregates as well as experimental advice. MN Waxham, KF Tolias, and SJ Ludtke offered conceptual guidance, and KF Tolias and SJ Ludtke supervised the project. All authors edited and reviewed the manuscript.

## References

1. Tsolaki E, Csincsik L, Xue J, Lengyel I, Bertazzo S. Nuclear and cellular, micro and nano calcification in Alzheimer’s disease patients and correlation to phosphorylated Tau. Acta Biomater. 2022;143:138–44.

2. Monfrini E, Arienti F, Rinchetti P, Lotti F, Riboldi GM. Brain Calcifications: Genetic, Molecular, and Clinical Aspects. Int J Mol Sci. NLM (Medline); 2023.

3. Loeb JA, Sohrab SA, Huq M, Fuerst DR. Brain calcifications induce neurological dysfunction that can be reversed by a bone drug. J Neurol Sci. 2006;243:77–81.

4. Termine JD, Posner AS. Calcium Phosphate Formation in vitro 1. Factors Affecting Initial Phase Separation’. Arch Biochem Biophys. 1970.

5. Mahamid J, Sharir A, Gur D, Zelzer E, Addadi L, Weiner S. Bone mineralization proceeds through intracellular calcium phosphate loaded vesicles: A cryo-electron microscopy study. J Struct Biol. 2011;174:527–35.

6. Mahamid J, Sharir A, Addadi L, Weiner S. Amorphous calcium phosphate is a major component of the forming fin bones of zebrafish: Indications for an amorphous precursor phase [Internet]. 2008. Available from: www.pnas.org/cgi/content/full/

7. Crane NJ, Popescu V, Morris MD, Steenhuis P, Ignelzi MA. Raman spectroscopic evidence for octacalcium phosphate and other transient mineral species deposited during intramembranous mineralization. Bone. 2006;39:434–42.

8. Hasegawa T, Yamamoto T, Tsuchiya E, Hongo H, Tsuboi K, Kudo A, et al. Ultrastructural and biochemical aspects of matrix vesicle-mediated mineralization. Japanese Dental Science Review. Elsevier Ltd; 2017. p. 34–45.

9. Thouverey C, Malinowska A, Balcerzak M, Strzelecka-Kiliszek A, Buchet R, Dadlez M, et al. Proteomic characterization of biogenesis and functions of matrix vesicles released from mineralizing human osteoblast-like cells. J Proteomics. 2011;74:1123–34.

10. Genge BR, Wu LNY, Wuthier RE. Differential fractionation of matrix vesicle proteins: Further characterization of the acidic phospholipid-dependent Ca2+-binding proteins. Journal of Biological Chemistry. 1990;265:4703–10.

11. Müller KH, Hayward R, Rajan R, Whitehead M, Cobb AM, Ahmad S, et al. Poly(ADP-Ribose) Links the DNA Damage Response and Biomineralization. Cell Rep. 2019;27:3124–3138.e13.

12. Greenawalt JW, Rossi CS, Lehninger AL. EFFECT OF ACTIVE ACCUMULATION OF CALCIUM AND PHOSPHATE IONS ON THE STRUCTURE OF RAT LIVER MITOCHONDRIA. J Cell Biol. 1964;23:21–38.

13. Boonrungsiman S, Gentleman E, Carzaniga R, Evans ND, McComb DW, Porter AE, et al. The role of intracellular calcium phosphate in osteoblast-mediated bone apatite formation. Proc Natl Acad Sci U S A. 2012;109:14170–5.

14. Pei D dan, Sun J long, Zhu C hui, Tian F cong, Jiao K, Anderson MR, et al. Contribution of Mitophagy to Cell-Mediated Mineralization: Revisiting a 50-Year-Old Conundrum. Advanced Science. 2018;5.

15. Wuthier RE, Rice GS, Wallace JEB, Weaver RL, Legeros RZ, David Eanes E. Calcified Tissue International Precipitation of Calcium Phosphate Under IntraceUular Conditions: Formation of Brushite from an Amorphous Precursor in the Absence of ATP. Calcif Tissue Int. 1985.

16. Duvvuri B, Lood C. Mitochondrial Calcification. Immunometabolism (United States). Lippincott Williams and Wilkins; 2021.

17. Barnard T, Ruusa J. Mitochondrial matrix granules in soft tissues. I. Elemental composition by x-ray microanalysis. Exp Cell Res. 1979;124:339–47.

18. Maetzler W, Stünitz H, Bendfeldt K, Vollenweider F, Schwaller B, Nitsch C. Microcalcification after excitotoxicity is enhanced in transgenic mice expressing parvalbumin in all neurones, may commence in neuronal mitochondria and undergoes structural modifications over time. Neuropathol Appl Neurobiol. 2009;35:165–77.

19. Jennings RB. Historical perspective on the pathology of myocardial ischemia/reperfusion injury. Circ Res. 2013. p. 428–38.

20. Halestrap AP, Clarke SJ, Javadov SA. Mitochondrial permeability transition pore opening during myocardial reperfusion - A target for cardioprotection. Cardiovasc Res. 2004. p. 372–85.

21. Rottenberg H, Hoek JB. The mitochondrial permeability transition: Nexus of aging, disease and longevity. Cells. MDPI; 2021. p. 1–23.

22. Jiao H, Jiang D, Hu X, Du W, Ji L, Yang Y, et al. Mitocytosis, a migrasome-mediated mitochondrial quality-control process. Cell. 2021;184:2896–2910.e13.

23. Ma L, Li Y, Peng J, Wu D, Zhao X, Cui Y, et al. Discovery of the migrasome, an organelle mediating release of cytoplasmic contents during cell migration. Cell Res. 2015;25:24–38.

24. Ma Y, Li T, Zhao L, Zhou D, Dong L, Xu Z, et al. Isolation and characterization of extracellular vesicle-like nanoparticles derived from migrasomes. FEBS Journal. 2023;290:3359–68.

25. Tolias KF, Bikoff JB, Burette A, Paradis S, Harrar D, Tavazoie S, et al. The Rac1-GEF Tiam1 couples the NMDA receptor to the activity-dependent development of dendritic arbors and spines. Neuron. 2005;45:525–38.

26. Duman JG, Tzeng CP, Tu YK, Munjal T, Schwechter B, Ho Szu-Yu T, et al. The adhesion-GPCR BAI1 regulates synaptogenesis by controlling the recruitment of the Par3/Tiam1 polarity complex to synaptic sites. Journal of Neuroscience. 2013;33:6974–8.

27. Pivovarova NB, Nguyen H V., Winters CA, Brantner CA, Smith CL, Andrews SB. Excitotoxic calcium overload in a subpopulation of mitochondria triggers delayed death in hippocampal neurons. Journal of Neuroscience. 2004;24:5611–22.

28. Lemieux H, Blier PU, Gnaiger E. Remodeling pathway control of mitochondrial respiratory capacity by temperature in mouse heart: Electron flow through the Q-junction in permeabilized fibers. Sci Rep. 2017;7.

29. Wu GH, Smith-Geater C, Galaz-Montoya JG, Gu Y, Gupte SR, Aviner R, et al. CryoET reveals organelle phenotypes in huntington disease patient iPSC-derived and mouse primary neurons. Nat Commun. 2023;14.

30. Fischer TD, Dash PK, Liu J, Waxham MN. Morphology of mitochondria in spatially restricted axons revealed by cryo-electron tomography. PLoS Biol. 2018;16.

31. Wilson LT, Tipping WJ, Wetherill C, Henley Z, Faulds K, Graham D, et al. Mitokyne: A Ratiometric Raman Probe for Mitochondrial pH. Anal Chem. 2021;93:12786–92.

32. Habraken WJEM, Tao J, Brylka LJ, Friedrich H, Bertinetti L, Schenk AS, et al. Ion-association complexes unite classical and non-classical theories for the biomimetic nucleation of calcium phosphate. Nat Commun. 2013;4.

33. Dowell LG, Rinfret AP. LOW-TEMPERATURE FORMS OF ICE AS STUDIED BY X-RAY DIFFRACTION. Nature. 1960;188:1144–8.

34. Uoselis L, Nguyen TN, Lazarou M. Mitochondrial degradation: Mitophagy and beyond. Mol Cell. Cell Press; 2023. p. 3404–20.

35. Zhao X, Lei Y, Zheng J, Peng J, Li Y, Yu L, et al. Identification of markers for migrasome detection. Cell Discov. Nature Publishing Groups; 2019.

36. Jasielec JJ, Filipek R, Dołowy K, Lewenstam A. Precipitation of inorganic salts in mitochondrial matrix. Membranes (Basel). 2020;10.

37. Tsolaki E, Csincsik L, Xue J, Lengyel I, Bertazzo S. Nuclear and cellular, micro and nano calcification in Alzheimer’s disease patients and correlation to phosphorylated Tau. Acta Biomater. 2022;143:138–44.

38. Schober R, Hilbrich I, Jäger C, Holzer M. Senile plaque calcification of the lamina circumvoluta medullaris in Alzheimer’s disease. Neuropathology. 2021;41:366–70.

39. Chen M, Bell JM, Shi X, Sun SY, Wang Z, Ludtke SJ. A complete data processing workflow for cryo-ET and subtomogram averaging. Nat Methods. 2019;16:1161–8.

40. Pettersen EF, Goddard TD, Huang CC, Meng EC, Couch GS, Croll TI, et al. UCSF ChimeraX: Structure visualization for researchers, educators, and developers. Protein Science. 2021;30:70–82.

41. Dang L, Ludtke SJ. Object Segmentation on Cryo-electron Tomography Data. Microscopy and Microanalysis. 2022;28:1514–6.

42. Tang G, Peng L, Baldwin PR, Mann DS, Jiang W, Rees I, et al. EMAN2: An extensible image processing suite for electron microscopy. J Struct Biol. 2007;157:38–46.

